# Gut bacteria-derived sphingolipids alter innate immune responses to oral cholera vaccine antigens

**DOI:** 10.1101/2021.12.01.470820

**Authors:** Denise Chac, Fred J. Heller, Hasan Al Banna, M. Hasanul Kaisar, Fahima Chowdhury, Taufiqur R. Bhuiyan, Afroza Akter, Ashraful I. Khan, Mia G. Dumayas, Susan M. Markiewicz, Amelia Rice, Polash Chandra Karmakar, Pinki Dash, Regina C. LaRocque, Edward T. Ryan, Samuel S. Minot, Jason B. Harris, Firdausi Qadri, Ana A. Weil

**Affiliations:** Department of Medicine, University of Washington, Seattle, WA, USA; Duke University School of Medicine, Durham, NC, USA; Infectious Diseases Division, International Centre for Diarrheoal Disease Research, Bangladesh, Dhaka, Bangladesh; Department of Medicine, Harvard Medical School, Boston, MA, USA; Department of Immunology and Infectious Diseases, Harvard School of Public Health, Boston, MA, USA; Data Core, Fred Hutchison Cancer Research Center, Seattle, WA, USA; Department of Pediatrics, Harvard Medical School, Boston, MA, USA; Division of Global Health, MassGeneral Hospital for Children, Boston, MA, USA; Department of Global Health, University of Washington, Seattle, WA, USA

**Keywords:** cholera vaccine, *Vibrio cholerae*, cholera, gut microbiome, microbiota, oral vaccine, sphingolipids

## Abstract

The degree of protection conferred after receiving an oral cholera vaccine (OCV) varies based on age, prior exposure to *Vibrio cholerae*, and unknown factors. Recent evidence suggests that the microbiota may mediate some of the unexplained differences in oral vaccine responses. We used metagenomic sequencing of the microbiota at the time of vaccination, and then related microbial features to immune responses after OCV using a reference-independent gene-level analysis. We found that the presence of sphingolipid-producing bacteria was associated with the development of protective immune responses after OCV. We experimentally tested these results by stimulating human macrophages with *Bacteroides xylanisolvens* metabolites and found that sphingolipid-containing extracts increased innate immune responses to OCV antigens. Our findings demonstrate a new analytic method for translating metagenomic sequencing data into strain-specific results associated with a biological outcome, and in validating this tool, we identified that microbe-derived sphingolipids impact *in vitro* immune responses to OCV antigens.

## INTRODUCTION

Cholera is a severe diarrheal disease caused by the bacterium *Vibrio cholerae*, and millions of cases occur annually.^1^ Despite increased surveillance and ongoing efforts to provide safe water to vulnerable populations, there has been recent re-emergence or new outbreaks of cholera in Haiti, Lebanon, and Syria, in addition to ongoing endemic spread in over 50 countries in Asia and Africa.^2–4^ Oral cholera vaccines (OCV) are important tools in cholera control but provide a short duration of protection and are less effective in children under five years of age.^5^

Long-term immunologic protection from cholera is incompletely understood. The vibriocidal titer is a commonly used measure of the humoral response to *V. cholerae* infection, and although higher titers are associated with protection from disease, titers wane prior to loss of protective immunity.^6,7^ Antibody responses to natural infection are largely directed toward cholera toxin (CT) and the *V. cholerae* lipopolysaccharide (LPS).^8^ In contrast with other toxin-mediated infections, CT-specific antibody responses are associated with only short-term protection from cholera.^5,9,10^ The *V. cholerae* O1 serogroup is responsible for the majority of epidemic cholera, and serogroup specificity is defined by the composition of the O-specific polysaccharide (OSP) of the LPS. After infection, OSP-specific memory B cells (MBCs) are detectable for up to one year^11^, and upon exposure to *V. cholerae,* these cells are presumed to differentiate rapidly into antibody-secreting plasma cells to produce antibodies that act at the mucosal surface to protect from infection.^9,12–15^ This is supported by longitudinal studies of household contacts of cholera patients, in which persons with elevated OSP-specific MBCs at the time of exposure to *V. cholerae* were protected from infection.^9,16^ Recent *in vivo* studies established that OSP-specific antibodies reduce *V. cholerae* colonization in a motility-dependent manner, and thus, are thought to be a mediator of protection in human infection.^17^ A collection of genetic, genomic, and laboratory-based studies have established that innate immune responses occur after acute infection and have an important role in shaping long term memory B cell responses.^12,15,18–20^ Innate immune responses contribute to the development and growth of OSP-specific MBCs that later reactivate by promoting upregulation of effector proteins and stimulating T follicular helper cell development for provision of “help” to B cells in germinal centers.^21–23^

On a population level, vaccination with a whole-cell killed OCV stimulates an OSP-specific MBC response in half of OCV recipients.^11,24–26^ While demographic factors such as age correlate with the likelihood of developing a protective vaccine response, biological mechanisms of these response differences have not been identified ^25,27,28^ The concept that gut microbiota differences may mediate differences in host response to OCV is supported by associations between the microbiota and responses to other oral vaccines,^29,30^ the observation that immune responses to OCV were reduced in recipients with bacterial intestinal overgrowth, and our prior study that detected associations between the gut microbiota and OSP-specific MBC responses to OCV in a separate cohort.^24,31–33^ To understand how gut microbes present at the time of vaccination could impact the host response to OCV, we applied a novel gene-level association method to metagenomic sequencing data from the microbiota at the time of OCV receipt. This analytic technique was selected and adapted to our analysis to identify specific microbial isolates associated with our outcome of interest, and this enabled us to experimentally test our sequencing study findings. We found that the presence of *Bacteroides* species capable of producing sphingolipids were correlated with the development of protective immune responses after OCV. To validate these findings, we exposed human macrophages to *Bacteroides*-derived sphingolipids *in vitro* and found a greater innate immune response to *V. cholerae* antigens, which suggests that gut microbial metabolites may impact immune responses to OCV in this population.

## RESULTS

### Immune responses to OCV are not correlated to simple microbiome composition

Eighty-nine participants received a whole-cell killed *V. cholerae* OCV with recombinant cholera toxin B (WR-rCTB) and underwent measurements of post-vaccine immune response in Dhaka, Bangladesh.^26^ Stool collected on the day of OCV administration underwent metagenomic sequencing. Twenty-two participants were aged 2-5 years (11/22, 50% female), 22 participants >5-17 years (12/22, 54% female), and 45 participants ≥18-46 years (32/45, 71% female). Immune responses after OCV were consistent with prior studies of whole-cell killed vaccine responses from populations living in cholera-endemic areas.^11,24,28,34^ As described in the parent study^26^, vibriocidal responses to *V. cholerae* O1 Ogawa and Inaba increased after vaccination, and some participants had an increase in MBC response, as observed in prior studies.^25,35^ Mean increases in CT- and OSP-specific IgG-producing MBCs before and after vaccination were significantly increased, while CT- or OSP-specific MBCs producing IgA were not, as previously seen.^26^ We found that OCV recipients were more likely to develop IgG directed OSP-specific MBC responses than IgA OSP-specific responses (**Figure 1A**). Among the immunologic parameters measured in this study, we focused primarily on *V. cholerae* OSP-specific MBC results because the presence of detectable OSP-specific MBCs is associated with long-term protection from infection, and a potential mechanism for OSP-specific antibody protective immunity has recently been established.^9,16,17^ To assess microbiome differences among vaccine recipients, vaccine responders (R) and nonresponders (NR) were categorized according to development of an MBC response between pre- and post-vaccination timepoints, or no change in MBC measures between timepoints, respectively.

**Figure 1.**
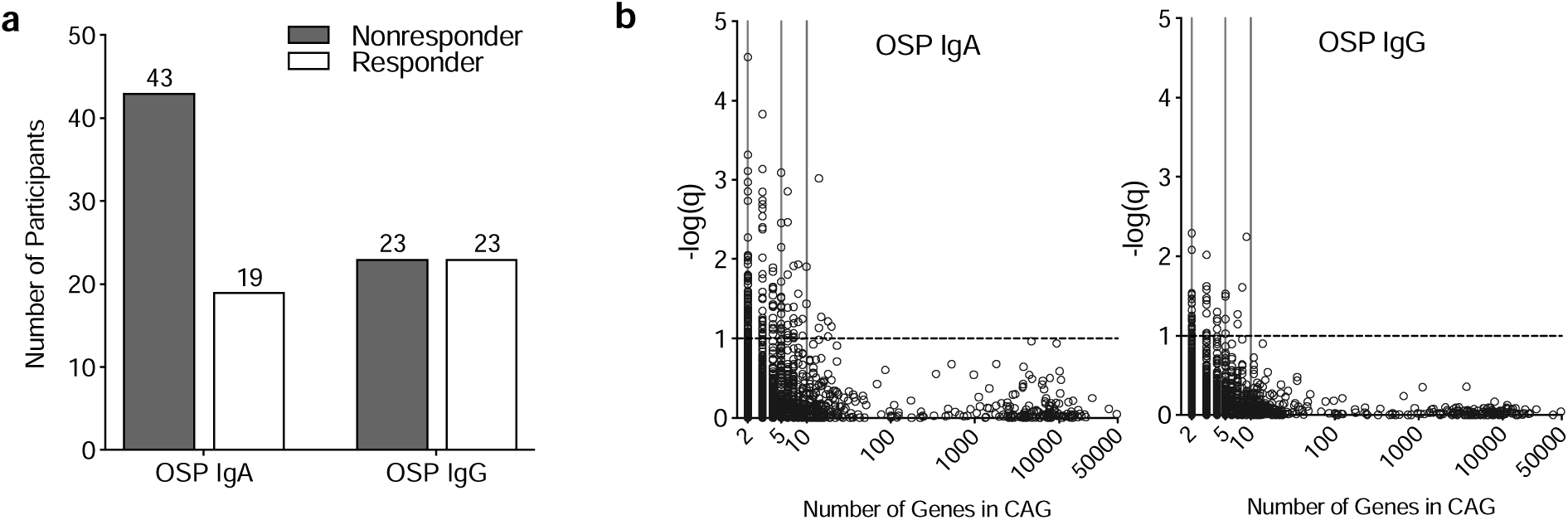
Co-abundant groups of genes in vaccine participant stool are associated with protective memory B-cell responses to OCV. (**a**) Counts of vaccine responder and nonresponder participants by OSP-associated MBC responses. Responders were defined as an OCV recipient who developed an increase in MBC response between pre- and post-vaccination timepoints, and nonresponders were defined as having no change in MBC responses between timepoints. (**b**) CAGs shown by size distribution and association with MBC response measures. CAGs are plotted as open circles based on the number of genes in a CAG along the x-axis and the -log of FDR adjusted q-value along the y-axis. The dotted line corresponds to q=0.1 and represents the pre-determined threshold for CAG significance.

We compared microbial taxonomic diversity measures by age and sex and found no significant differences by age (**Figure S1A-B**). Shannon diversity (a measure of both species diversity and abundance) demonstrated a difference in microbial diversity by sex (**Figure S1C-E**), driven by fewer *Lactobacillales, Veillonellaceae,* and *Eubacteriaceae,* and more *Erysipelotrichales, Ruminococcaceae, and Lachnospiraceae* in women, consistent with microbiome-specific sex differences noted in other populations.^36^ No significant difference was found by Shannon Index or by Bray-Curtis community structure between vaccine responders and nonresponders for the four antigen-specific MBC measures (IgG and IgA producing OSP-specific or CT-specific MBCs, **Figure S2A-B**).

### Co-abundant gene groups differentiate the microbiota of OCV responders from nonresponders

To address the high dimensionality of the metagenomic data and examine detailed features of the microbiota, we used gene groups as the fundamental unit of analysis. Co-abundant genes (CAGs) are likely to be encoded on a shared genetic element (e.g. chromosome or plasmid) across the organisms present in the cohort, and grouping genes together in this way has been successfully used to infer the relationship between specific genes and biological outcomes.^37–39^ We identified CAGs which were associated with MBC responses and predetermined a significance threshold of a False-Discovery Rate (FDR)-adjusted q-value of ≤0.1, based on prior studies associating metagenomic data with biological responses (**Figure 1B**).^40^ We calculated significant CAGs for each vaccine response measure separately (IgA OSP, IgA CT, IgG OSP, and IgG CT), and identified CAGs associated with an MBC response or lack of MBC response in each group. A total of 4147 CAGs were identified, and 323 CAGs containing ≥2 genes met our significance threshold across the four vaccine response measures (**Table S1**). The greatest number of significant CAGs that mapped to bacterial genomes were associated with IgA OSP-specific MBC responses, and most of these CAGs were associated with a higher immune response (**Table S1**). CAGs mapped to assembled bacterial genomes demonstrated that many CAGs aligned to multiple species (**Figure S3**). Many genes composing significant CAGs have unknown functions or hypothetical proteins (based on the results of the eggNOG functional annotation and NCBI nr/blastp database **Table S2**). Because all human sequences were removed from metagenomic data prior to CAG construction, this represents an enormous underexplored and unidentified microbial diversity of possible clinical significance that may be bacterial, fungal, or viral in origin. Among annotated genes in significant CAGs, most encoded nonspecific cellular functions, were related to amino acid metabolism, or were phage associated.

### Co-abundant gene group association with OCV responses were strain specific

Because CAGs mapped to multiple reference strains, a priority score was assigned to each bacterial strain detected from participants in this study (see Methods). This process resulted in top bacterial strains associated with each MBC response, and many were from diverse taxonomic groups (**Figure 2A-B, Figure S4A-B, Table S3,** strain listing with priority scores **Table S4-7**). Typically, only a small fraction of strains within a given taxonomic group had a high priority score, indicating that the association between MBC responses and gut microbes may be strain specific (**Figure 2C, Figure S4C**). Among 412 total bacterial genera detected in the stool of the study population, only 34 contained five or more strains that mapped to a significant CAG (**Figure 2C, Figure S4C**). To demonstrate the relationship between high priority and unscored strains within one species, we first identified *Bacteroides xylanisolvens* strains that were detected in the stool of our study population. We next created a phylogenetic tree using NCBI genomes of all *Bacteroides xylanisolvens* strains (**Figure S5**). The four most highly ranked *B. xylanisolvens* strains were not consistently clustered (**Figure 2D**), suggesting that even closely related strains differ enough in gene content to be associated with different biological effects and is not restricted to a single monophyletic lineage or clade. One exception to this finding is that most strains of *Clostridioides difficile* strains detected in our study population were highly positively associated with this MBC response (**Figure 2C**). Among the 1,080 strains identified in study participant stool that positively correlated with OSP IgA MBC responses, *Bacteroides* species (n=31) and *Prevotella copri* (n=53) were disproportionately represented among the highest priority scored strains (**Figure 2B, Table S4**). Strains positively associated with OSP IgG responses were taxonomically varied and included members of *Bifidobacterium* and *Ruminococcus/Blautia* genera and several Enterobacteriaceae (including *Escherichia coli, Enterobacter,* and *Klebsiella* species) (**Figure 2A**, **Table S5**). Our results indicate that associations with biological outcomes were strain specific, and thus, strain-specific methods of identifying these relationships are needed to select appropriate strains for experiments to test and validate a correlative host-microbe interaction. These findings support the concept that taxonomic groupings alone are unreliable markers for specific genes or associations with biological effects.

**Figure 2.**
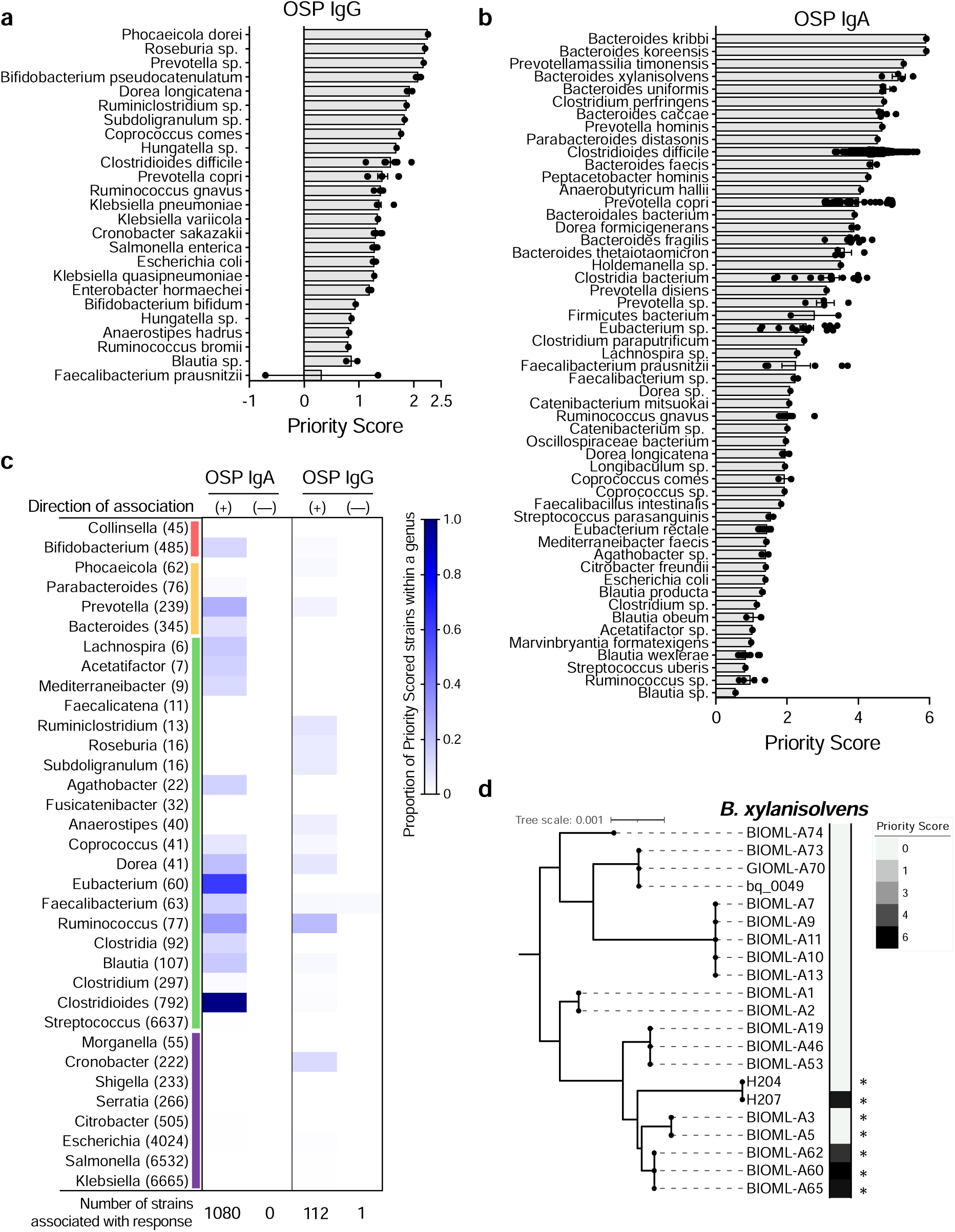
Co-abundant gene groups associated with OSP-specific memory B cell responses are strain-specific. CAGs were mapped onto all complete NCBI bacterial genomes as described in the methods, and species represented by significant CAGs with a priority score greater than 0.5 are shown for (**a**) OSP IgG and (**b)** OSP IgA MBC responses. Priority scores quantify the association between these strains and their association with a specific MBC response (see Methods). Each dot indicates the priority score of a specific strain within the species in the row. Bar represents mean with SEM of priority scored strains. (**c**) Proportion of strains found in our study population within each genus that were associated with a specific MBC response with positive (+) and (-) associations. Genera that contained 5 or more strains with significant CAGs were plotted with shading representing the percentage of total strains that contained significant CAGs. Parentheses indicate the total number of distinct strains in each genus found in our study population. Phylum identifications are shown in the colored vertical bar: red=Bacteroides; orange=Actinobacteria; green=Firmicutes; purple=Proteobacteria. For example, approximately 15% of the 485 Bifidobacterium strains detected in the study population were assigned a positive priority score associated with OSP IgA responses. (**d**) Priority scores differ widely among closely related strains within one species. Phylogenetic tree of Bacteroides xylanisolvens strains found in our study population and their priority scores representing their association with the OSP IgA MBC response. Tree made using NCBI published genomes of *Bacteroides xylanisolvens*. Asterisks (*) indicate presence of serine palmitoyl transferase (*spt*) orthologs. Spt is the enzyme required for bacteria to de novo synthesize sphingolipids.

### The microbiota of vaccine responders was characterized by bacterial species with sphingolipid producing potential

The identification of bacterial strains associated with OSP-specific IgA-producing MBC responses featured genera with the ability to produce sphingolipids, such as *Bacteroides* and *Prevotella* (**Figure 2B-D**). This was a notable finding because bacteria rarely produce their own sphingolipids, and these metabolites are often unique, odd-chain lipid structures that are distinct from eukaryotic sphingolipids.^41–43^ Recent studies have demonstrated that *Bacteroides-*derived sphingolipids reduce intestinal mucosal inflammation.^34,35^ For these reasons, we investigated the relationship between vaccine response and *Bacteroides*-derived lipids. First, we characterized total lipid content in study participant stool using targeted, quantitative mass spectrometry-based lipidomics. Several sphingolipid derivatives, including ceramides, were more abundant among vaccine responders (**Figure 3A** and **Table S8**). While whole stool lipidomics did not specifically distinguish between host- or bacterial-derived lipids, we were interested in these differences in lipid content between OCV responders and nonresponders because in our prior study, fecal extracts from these two groups generated different innate immune responses in our model of OCV response.^24^ We next used the metagenomic data to assess whether vaccine recipient stool differed in serine palmitoyltransferase (*spt*) gene abundance, the enzyme required for *de novo* assembly of microbe-derived sphingolipids.^44^ We found that vaccine responders had higher abundance of *Bacteroides spt* compared to nonresponders (**Figure 3B**).

**Figure 3.**
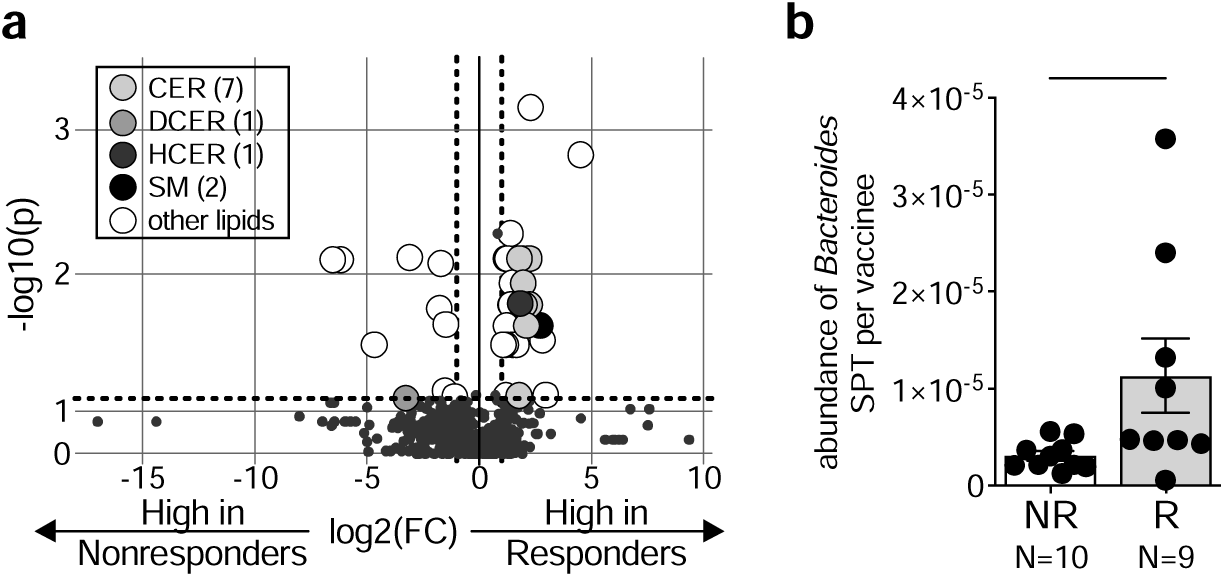
Bacteroides *serine palmitoyltransferase spt* gene presence in stool is associated with a protective memory B cell response to OCV. (**a**) Targeted quantitative mass spectrometry based lipidomics of fecal samples from vaccine responders and nonresponders at the time OCV was administered. P values from Multiple Mann-Whitney tests are displayed on the y-axis with -log10 transformation and x-axis is fold change with log2 transformation. Dotted lines represent fold change of 1 or -1, and P-value of 0.05 along the x- and y-axis, respectfully. (**b**) Relative abundance of *Bacteroides spt* among 19 select vaccine responders (R) who had both an IgA and IgG-specific OSP response to OCV and nonresponders (NR) who lacked both of these OSP responses. Each dot represents one person. Bars represent mean with SEM. Mann-Whitney test was performed; *,P< 0.05. CER: ceramides, DCER: dihydroceramides, HCER: hexosylceramides, SM: sphingomyelin.

### *Bacteroides*-produced sphingolipids blunted inflammatory cytokine production in human macrophages

Since innate immune responses impact the development of subsequent adaptive immune responses including the differentiation and proliferation of MBCs, we next tested whether the production of *Bacteroides* sphingolipids impacted mucosal innate immune responses in our human macrophage cell culture model of OCV response.^24,45^ Since multiple strains of *Bacteroides xylanisolvens* (Bx) were associated with OSP-IgA MBC responses to OCV, we selected an *spt*-positive Bx strain for these additional experiments. This strain was isolated from the stool of one of our study participants. We exposed THP-1-derived macrophages to lysate from Bx grown with or without the presence of myriocin, an *spt* inhibitor^46^, and measured cytokine responses. Myriocin-treated, sphingolipid-free (SL-free) Bx lysate induced significantly more interleukin-6 (IL-6) compared to the untreated, sphingolipid-containing (SL-containing) lysate when applied to THP-1 macrophages (**Figure S6**). To identify which component of the bacterial lysate contributed to this difference in response, Bx lysates were fractionated into culture supernatant metabolites, non-soluble cell membrane, and lipid components using the Bligh-Dyer method.^47^ Metabolite and non-soluble fractions of culture supernatant did not generate different cytokine responses between SL-free and SL-containing fractions, but increased IL-6, tumor necrosis factor-*α* (TNF-*α*), and IL-8 were produced when SL-free lipid fractions were used, compared to SL-containing lipid fractions (**Figure 4A-C**). The production of IL-6 in wells with SL-free Bx lipids was nearly 6-fold less than wells exposed to Bx sphingolipids.

**Figure 4.**
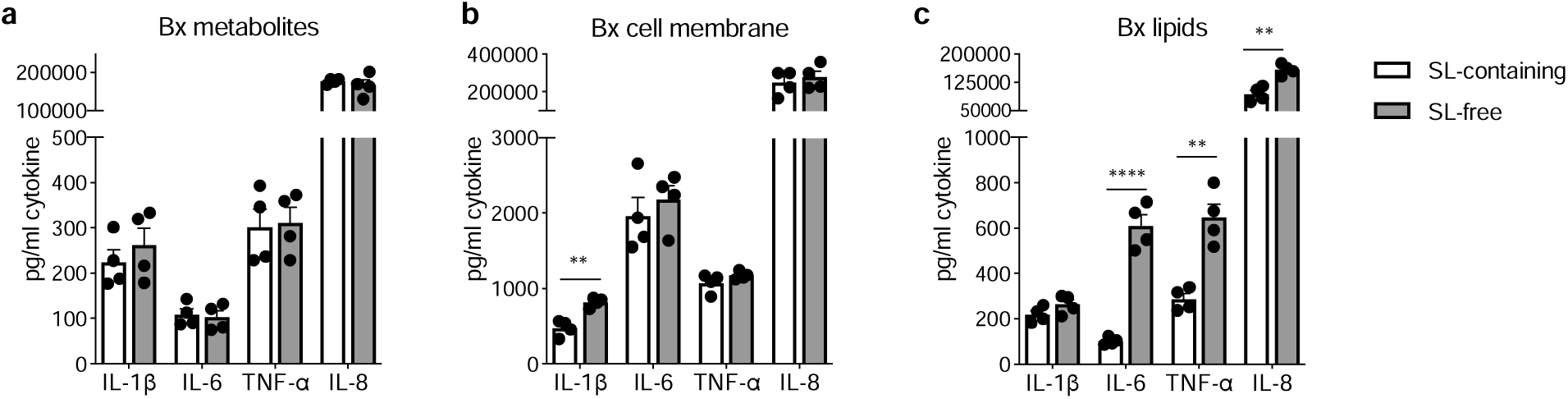
Exposure to *B. xylanisolvens*-derived sphingolipids reduces immune activation of human THP-1 macrophages. A *B. xylanisolvens* isolate from human stool was cultured with or without myriocin (SL-free and SL-containing, respectively) and the bacterial lysate was fractionated using Bligh Dyer lipid extraction. Cytokine responses were measured in supernatant of THP1-derived macrophages treated for 18 hours with B. xylanisolvens (**a**) metabolites, (**b**) non-soluble cell membrane components, and (**c**) lipids. ELISAs were used to measure cytokine response. Statistical analysis was performed using multiple unpaired t tests with FDR two-stage set-up method of Benjamini, Kriege and Yekutieli. **, P<0.01; ****, P<0.0001. Bars indicate mean with SEM.

### *Bacteroides*-produced sphingolipids increased innate immune responses to *V. cholerae* antigens

We observed an increase in innate cytokine responses when Bx SL-free lipid extracts were applied to human macrophages, and this was consistent with our prior findings that fecal extracts from vaccine nonresponders increased IL-6 production in this model.^24^ Because this response was triggered by vaccine nonresponders only, we hypothesized that an increase in resting innate immune activation could dampen the immune response to vaccine antigens. To test this, we pretreated THP-1-derived macrophages with SL-free and SL-containing lipid fractions and then exposed them to heat-killed *V. cholerae* strain JBK70, an A-B toxin negative vaccine strain, as a proxy for OCV, as in prior studies.^45^ In contrast to the decreased inflammatory cytokine response observed in our model after exposure to SL-containing Bx supernatant, macrophages that were pretreated with SL-containing lipid fractions and then stimulated with JBK70 produced increased IL-6, monocyte chemoattractant protein-1 (MCP-1), and TNF-*α* (**Figure 5A**). We next removed phospholipids from all lipid preparations using a mild alkaline hydrolysis treatment.^46^ Alkaline treatment of SL-containing lipid fractions further increased the magnitude of IL-1*β*, IL-6, IL-12p40, and TNF-*α* production in response to JBK70 stimulation (**Figure S7**). To test for this effect in another *spt-*positive *Bacteroides* isolate from our study population, we selected a *Bacteroides koreensis*, a species identified as highly associated with OSP-specific MBC OCV responses. Our findings were reliably replicated using this strain (**Figure 2B, S8**).

**Figure 5.**
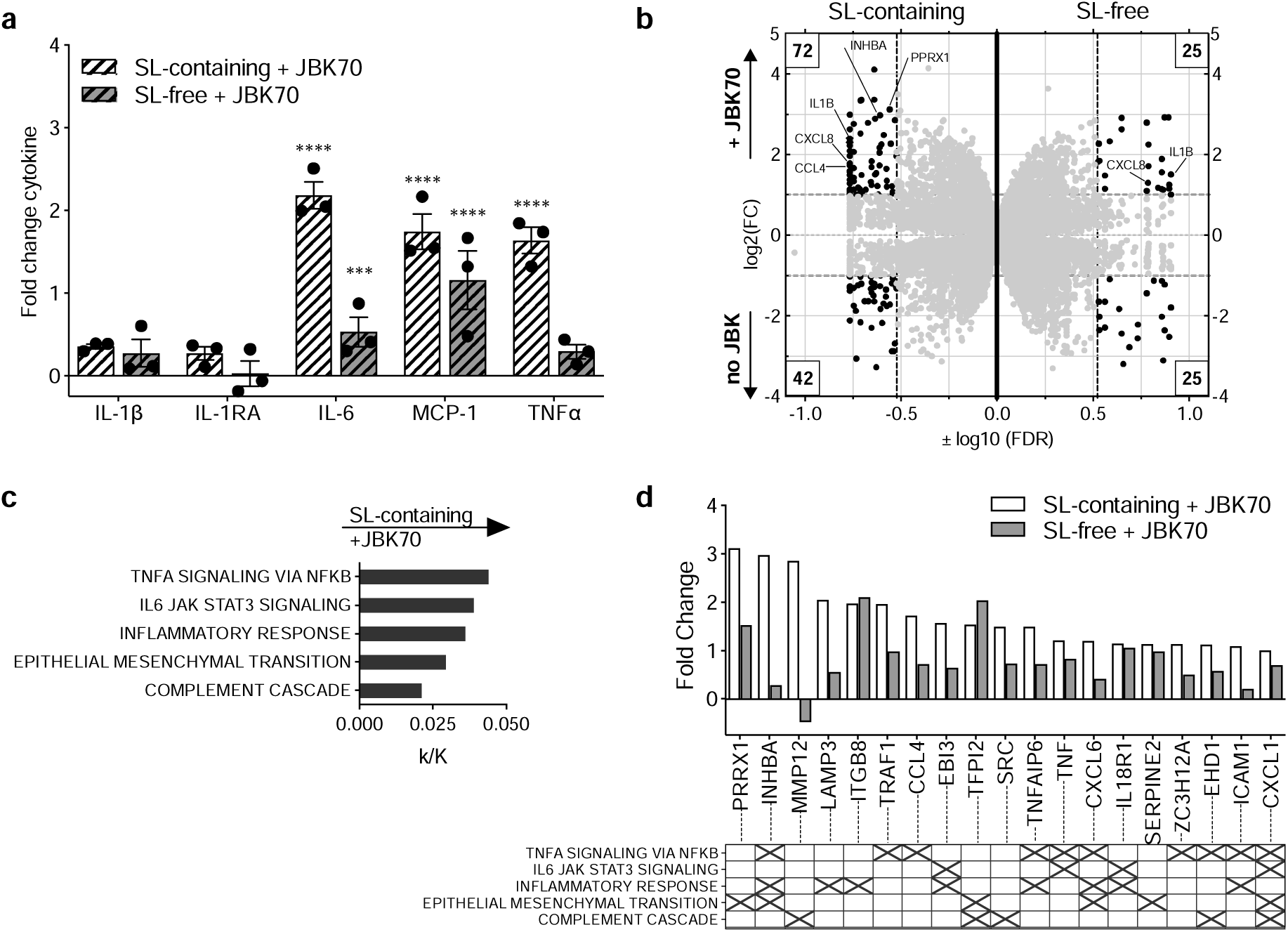
*B. xylanisolvens*-derived sphingolipids increase inflammatory cytokine responses from THP-1 human macrophages after stimulation with a *V. cholerae* vaccine strain. (**a**) Cytokine responses shown as fold change measured in supernatant from THP1-derived macrophages following preconditioning with Bx lipid extracts and treated with heat-killed JBK70. 2-way ANOVA on cytokine pg/mL with bars indicating mean with SEM is shown. ***, P<0.001; ****, P<0.0001. (**b**) Gene expression from THP1-derived macrophages measured by RNA sequencing following pre-conditioning with Bx SL-containing or SL-free lipid fractions with or without exposure to HK JBK70. X-axis is log10(FDR) and Y-axis is log2(fold change). Each gene is represented as one dot and is shown twice in this figure, once in the SL-containing (left side) that was preconditioned with sphingolipids and once in the SL free (right side) that was preconditioned without sphingolipids present. Numbers boxed in the corners indicate the number of genes with FDR < 0.3 and FC > 2. (**c**) Pathways enriched based on macrophage gene expression after treatment with SL-containing lipid fractions and JBK70 stimulation, compared to macrophages treated with SL-free lipid fractions, FDR values < 0.05. X axis label is proportion enriched (k/K) represented as ratio of significant genes in that pathway over total genes in the specified pathway, calculated using BIGprofiler through clusterProfile. (**d**) Fold change of genes identified in pathways enriched following SL-containing lipid fraction exposure and JBK70 treatment in THP-1 macrophages. Matrix below graph indicates gene membership in each pathway based on hallmark gene sets in the Molecular Signatures Database.

We next performed RNA-sequencing on RNA from THP-1 macrophages preconditioned with Bx SL-containing or SL-free lipid fractions with and without exposure to heat-killed JBK70 (**Figure 5B)**. Using a Gene Set Enrichment Analysis, we found preconditioning macrophages with SL-free Bx lipids prior to JBK70 exposure increased activation of several inflammatory pathways compared to cells preconditioned with SL-containing lipids (**Figure S9A**). Compared to the SL-containing fraction, SL-free fraction preconditioning triggered pathways associated with interferon-*α*, interferon-□, and TNF-*α* cytokine responses, and IL-6 – Janus kinase (JAK) – JAK-signal transducer and activator of transcription (STAT3) pathways. After preconditioning and subsequent exposure to heat-killed JBK70, TNF-*α* signaling via the nuclear factor kappa-light-chain-enhancer of B cells (NFκB) pathways were upregulated independent of SL exposure (**Figure S9B-C,** individual genes shown in **Table S9**).

To identify immune activation pathways that distinguished SL-free from SL-containing lipid stimulation of human macrophages after JBK-70 stimulation, we next excluded gene expression that occurred in response to heat-killed JBK70 independent of bacterial SL exposure. Here, human macrophage responses after SL-containing lipid treatment and JBK-70 stimulation were enriched in TNF-*α* signaling via NF-κB, IL-6 – JAK – STAT3 signaling, inflammatory response, epithelial/mesenchymal transition, and complement pathways, compared to macrophages treated with SL-free lipids (**Figure 5C**). The 19 genes differentially associated with enriched pathways are shown in **Figure 5D and Figure S10**. Several of the genes were associated with multiple inflammatory pathways including inhibin A (INHBA), C-X-C motif chemokine ligand 7 (CXCL7), and CXCL1. Overall, we found that human macrophage responses to JBK70 stimulation, a *V. cholerae* vaccine strain, are higher amplitude after exposure to *B. xylanisolvens* SL-containing lipid fractions.

## DISCUSSION

Oral cholera vaccines are an essential tool for reducing the global impact of cholera. Yet, the protection they provide is limited, and differs between individuals for unclear reasons. Improving the duration and degree of protection afforded by currently available OCVs would be a critical advance in cholera control. We previously found that the microbiome at the time of OCV administration was associated with the long-term MBC responses that are thought to mediate protection from cholera.^24^ To explore the relationship between the gut microbiota and host immune responses to OCV, we used whole genome sequencing (WGS) of the stool microbiota and applied a new analytic method to identify strain-specific correlations. We detected an association between commensal gut microbes that produce bacterial sphingolipids in stool from persons that developed protective immune responses after OCV, and *in vitro* experiments supported this finding and demonstrated greater innate immune responses to heat-killed JBK70, a *V. cholerae* vaccine strain. Taken together, these results suggest that gut microbial sphingolipids could enhance innate immune responses to oral cholera vaccines. Increasing the immunogenicity of existing cholera vaccine candidates that currently make up the World Health Organization vaccine stockpile would be a critical advance in the context of current vaccine shortages and increasing cholera outbreaks worldwide.

Previous studies of the relationship between the gut microbiota and responses to oral vaccination used sequencing methods that did not distinguish metagenomic content well enough to identify strain-level associations with vaccine response.^29,30,48,49^ Here, we describe a quantitative method of analyzing metagenomic sequence data that enables identification of bacterial strains associated with a biological outcome without dependence on a reference genome database or taxonomy for results interpretation. Because interpreting biological significance of *de novo* assembled gene content has serious limitations, we followed the CAG-level outcome association with exhaustive alignment of gene content against a collection of reference genomes. In this manner, we identified patterns of association tied to gene content which were orthogonal to a strict taxonomic hierarchy of microbial diversity. Supporting the value of our taxonomy-independent approach in this analysis, we observed no differences in alpha or beta diversity between vaccine responders and nonresponders for any of the vaccine response measures, and we found that strain-specific associations were not generalizable by genera or species.

To experimentally test our findings, we examined the relationship between sphingolipid-producing microbes that were associated with OSP-specific MBC responses to vaccination. Sphingolipid-producing microbes are generally represented in the human gut microbiota by two prominent commensal genera, *Prevotella* and *Bacteroides*.^50^ Microbe-derived sphingolipids from *Bacteroides* were previously observed to impact mucosal immune responses; for instance, Brown and colleagues found that the lack of *B. thetaiotaomicron* sphingolipids in mono-colonized mice resulted in increased IL-6 and MCP-1 produced *ex vivo* from epithelial cells.^44^ We identified a similar pattern of increased IL-6 and MCP-1 production in our model of OCV responses using human macrophages exposed to Bx sphingolipid-free lipid fractions. Based on the stimulatory response to Bx lipids that we observed in human macrophages, we were surprised to observe that exposure to sphingolipid-free Bx lipid fractions resulted in lower innate responses to a heat-killed *V. cholerae* strain used as a proxy for vaccine antigen. This inverse relationship, wherein higher baseline innate immune activation is correlated with a dampened immune response to vaccine antigen, has been previously described.^51–53^ For example, higher systemic innate immune responses at the time of Hepatitis B virus (HBV) vaccine administration correlated with lower peripheral immune responses in humans.^54,55^ HBV vaccine nonresponders were found to have much higher pre-vaccination gene expression of innate immune effectors and proteins including immune responses to LPS and defensins that activate NFκB signaling pathways.^56^ This effect has also been observed in human yellow fever vaccination, in which systemic measures of lymphocyte activation and high levels of proinflammatory monocytes at baseline negatively correlated with neutralizing antibody titers post-vaccination.^57^ This relationship has not been previously observed in oral vaccination, and warrants further exploration as a potential tool in improving responsiveness to OCVs and other oral vaccines. Although more testing is needed, the approach of using commensal gut microbes to condition or “prime” the mucosal surface for optimal responsiveness to vaccine antigens is novel and distinct from the use of traditional adjuvants that act by causing increased inflammation at the site of antigen delivery.

We measured the impact of microbial sphingolipids in our model of human innate immune responses to OCV because these responses support the differentiation and maintenance of antigen-specific B cells that mediate long-term protection from cholera. Sphingolipid-producing microbes were prominent among microbes associated with increased OCV responses in our analysis, and the plausibility of this relationship is supported by evidence from animal studies in which the presence of microbial sphingolipids was associated with more robust adaptive mucosal immune responses^58,59^. When we exposed human macrophages to a heat-killed OCV strain after pre-conditioning the cells with Bx sphingolipid-containing lipid fractions, we observed activation of several innate pathways characterized by genes typically expressed in bacterial infection and bacterial clearance^60–67^, and this was accompanied by expression of genes involved in inflammatory cell recruitment including leukocytes (via ICAM1), neutrophils (via CXCL1 and CXCL6), and dendritic cells and T-cells (via C-C motif chemokine ligand 4 (CCL4)).^60,68–70^ CCL4 is associated with the migration of CD4+ T cells and enhanced mucosal secretion of antigen-specific IgAs.^71–73^ This pattern of innate immune activation was significantly reduced when these lipids were absent at the time of OCV introduction, and downstream studies of these effects on other cell types are needed, in addition to the identification of the microbe-derived lipid species that produce this response.

Several co-abundant gene groups that were highly associated with immune responses to OCVs in our study mapped to pathogens, such as strains in the *Shigella* genus and strains of *Salmonella enterica*. Persons with diarrhea were excluded from this study, and these pathogens might have been detected in the stool due to subclinical or recent infection, or could be a marker of exposure to contaminated food and water, as has been described in other study participants living in areas lacking modern sanitation.^74–78^ Notably, we found that many strains of *Clostridioides difficile*, which can be a colonizing microbe in this population, were highly associated with IgA-specific OSP responses, but not other immune responses.^74^ Two *C. difficile* peptides (the fragment of the receptor-binding domain of toxin A TxA[C314] and a fragment of the 36 kDa surface-layer protein [SLP-36kDa] from strain C253) are known to have immunoadjuvant properties; investigators previously noted an increase in mucosal IgA and systemic IgG targeting antigen when these peptides were co-administered to mucosal surfaces in mice.^79^ This mechanism is a potential explanation for the strong correlation observed in our study between *C. difficile* and OSP IgA responses, and the specific pathways activated in response to OCV antigen will require further study. As expected, we found differences in the gut microbiota between males and females in this study; this was independent of overall immune responses to vaccination, and CAG distribution did not differ between sexes. We also did not collect data on antibiotic use or helminth infection, which are both widespread in this population, and could potentially impact immune responses.^80,81^ We were not able to control for the presence or absence of helminths in the study participant microbiota using sequencing data due to the limitations of metagenomic analysis including a relatively low abundance of eukaryotic reads compared to bacterial reads and a lack of accurate reference helminth genomes.^82,83^

In conclusion, we used a reference-independent gene-level metagenomic analysis of the gut microbiota at the time of vaccination to pinpoint specific strains that may govern the relationship between gut microbes and OCV responsiveness. Our findings strengthen the concept that individual differences in the microbiota drive variation in OCV response. Modification of baseline innate immune activation mediated by gut microbes could be a new target for improving OCV responses, such as through the use of pre- or probiotic products to be co-administered with OCVs.

## METHODS

### Study design and subject enrollment

Adult and child participants over 2 years of age and <60 years of age living in urban Dhaka, Bangladesh were enrolled through the International Centre for Diarrhoeal Disease Research, Bangladesh (icddr,b) if they had not previously been vaccinated with an OCV and had no major comorbid conditions.^84^ After obtaining written informed consent from patients or guardians, study participants received two doses of WC-rCTB doses (Dukoral), per the package insert, on study Day 0 and Day 14. Blood specimens were collected from the participants on Day -14 (two weeks prior to the first dose of vaccination), Day 2, Day 7, and Day 44. All timepoints are considered in reference to Day 0, the day of administration of the first dose. Vibriocidal titer was assessed at each of these timepoints; OSP-specific and CT-specific MBC were measured at Day -14 and Day 44. Demographic data were collected at enrollment and stool samples were collected at baseline prior to the first dose of WC-rCTB. Additional details and methods for vibriocidal titers, serum antibodies, and MBC measures are described in the parent study.^26^

### Stool processing and DNA preparation

Stool samples were selected for sequencing if the participant had MBC ELISPOT results for both pre-and post-vaccination timepoints for at least two out of four MBC responses. Application of this criteria resulted in sequencing the stool from 90% of study participants. Stool was collected in cryovials and immediately frozen at −80°C and shipped to US investigators. At the University of Washington, stool microbial DNA was extracted using Powersoil DNA isolation kit (Qiagen, Germany) with a modified protocol as previously described.^74^ Briefly, stool was thawed on ice and approximately 100 mg of stool was added to bead beating tubes. The stool was treated with C1 solution, heated at 65°C for 10 minutes, 95°C for 3 minutes, and vortexed for 10 minutes. Samples were treated using C2–C5 solutions provided by the Powersoil kit. Resulting DNA was eluted in DNase- and RNase-free water; quality and quantity were measured using a NanoDrop ND-1000 (Thermo Scientific, USA).

### Sequencing, read processing, and co-abundant gene (CAG) grouping

Sequencing was performed on the NovaSeq 6000 PE150 platform using standard Illumina adapters targeting a minimum of 10M reads per sample. The raw data can be accessed through NCBI, accession #PRJNA782606. Fastq files were processed using *geneshot*, a pipeline developed to streamline the analysis of gene-level metagenomic data and its association with experimental metadata.^37^ Using default settings for this pipeline (v0.9), raw WGS data from each individual specimen was quality-filtered and *de novo* assembled using Megahit^85^; human reads were excluded using human reference genome GRCh37 hg19, and protein-coding sequences were predicted using Prodigal^86^ and deduplicated across all specimens using linclust (at 90% amino acid similarity); coding sequences were annotated with taxonomic labels using DIAMOND (by alignment against NCBI RefSeq 01/22/2020) and with functional labels using eggNOG-mapper v2 (accessed 05/22/2024)^87^ and NCBI blastp (accessed 05/22/2024/05/22 with database nr_clustered(experimental)); WGS reads from each specimen were aligned against the *de novo* generated gene catalog with DIAMOND; coding sequences were grouped by co-abundance into CAGs.^38^ The centroid was selected for each deduplicated gene using the mmseqs2 algorithm.^88^ The depth of sequencing used for CAG generation was an average of the depth of sequencing across all genes in each CAG. The relationship between CAGs containing ≥2 genes were correlated with participant vaccine response measures using the corncob algorithm implementation of beta-binomial regression and accounting for multiple comparisons using the Benjamini-Hochberg false-discovery rate correction procedure.^89,90^ CAGs containing only one gene were excluded to reduce the total number of tested hypotheses, considering the relative lack of biological interpretability for singleton coding sequences in the context of gene-level metagenomic analysis. “Top” CAGs were defined prior to the analysis as those with a q-value ≤0.1.

### Assessment of gut microbial communities

Reads from all participant samples were used to estimate the relative abundance of taxonomic groups using metaphlan2^91^, which uses the alignment of raw reads against a set of previously identified marker genes that are conserved across taxonomic groupings of genomes. Output of metaphlan2 was merged across all samples and non-bacterial taxa were removed. Alpha diversity measures by Shannon Index and beta diversity measures by Bray-Curtis Dissimilarity Index were calculated using divnet (v3.6) in R (v4.0.2).^92^ Principal coordinate of analysis (PCoA) plots to visualize these results were generated using the R package vegan.

### CAG alignment to bacterial genomes

Top CAGs found to be significantly associated with vaccine response measures were aligned to bacterial genomes to identify the strains containing the most top CAGs. A total of 61,918 bacterial reference genomes were selected from the National Center for Biotechnology Information prokaryote genome database using DIAMOND v0.9.10 via the AMGMA (Annotation of Microbial Genomes By Microbiome Association) workflow (https://www.ncbi.nlm.nih.gov/genome/browse#!/prokaryotes/ and https://github.com/fredhutch/amgma).^38,93,94^ We used all bacterial sequences in the database apart from bacterial sequences with low assembly quality and completeness (chromosomes and scaffolds). We included all complete genomes and 85% of contig-level assemblies and excluded 15% of contig-level assemblies with the lowest levels of completeness (>200 contigs). Using the alignment of coding sequences from the metagenomic analysis against this set of partial and full genomes, a relative abundance measure was generated for each genome in each sample by summing the proportion of reads from a given sample that aligned to those genes found in each genome. Using that strain-level relative abundance measure, we applied the same statistical approach (beta-binomial modeling using the corncob algorithm) to estimate the coefficient of association with vaccine response measures on a per-genome basis.

### Development of CAG-based strain identification method

Once CAGs were aligned to bacterial genomes,^37,38^ we next quantified the association between these strains and vaccine response measures. For this purpose, we created a priority score designed to 1) account for the strength of the association between top CAGs aligning to a genome and the vaccine response measure of interest, 2) preserve the directionality of the association between the vaccine response measure and top CAGs, 3) be independent of CAG size, in order to avoid bias toward large CAGs containing more genes that could potentially align, and 4) discriminate between strains within the same species.

For a given strain with *n* number of significant CAGs aligning to strain genomes, this priority score was calculated as follows:

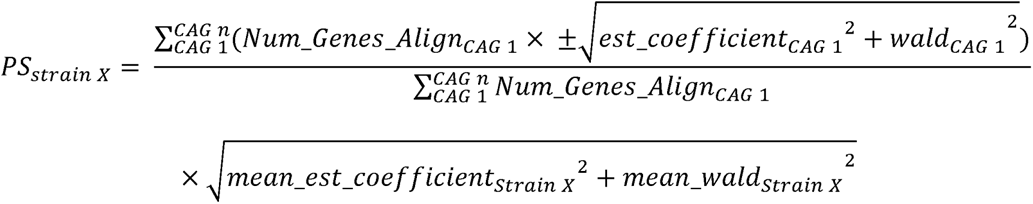

The estimated (est) coefficient reflects the abundance of a CAG in responders compared to nonresponders for each vaccine response measure, and the Wald score integrates standard error which reflects the uniformity of the association.^38^ The “±” in the first term corresponds to the sign of the CAG’s estimated coefficient. The number of genes aligning from a CAG to the strain genome is represented in the estimated coefficient and Wald score, creating a weighted measure of CAG-strain association which is summed in the first term’s numerator for all top CAGs with genes aligning to the strain. By normalizing this weighted sum in the numerator of the first term by the unweighted number of genes aligning from top CAGs, we minimize bias based on CAG size. In the second term, the score is further weighted by integrating the number of genes aligned to reference strains without regard for CAG grouping, and thereby discriminates between strains that may have otherwise identical first terms due to the alignment of the same number of genes from the same top CAG(s). A “+” priority score indicates a positive association between the vaccine response measure and strain, and a “-“ indicates an inverse relationship. The scripts used for priority score can be found in GitHub: https://github.com/letsgetthisfred/Dukoral_gene_level_analysis/blob/main/Generic%20Code%20for%20priority%20score%20and%20geneshot%20output.R).

### *spt* gene quantification

*Bacteroides* serine palmitoyltransferase (*spt*) abundance was determined by querying BT_0780, a confirmed spt gene in *Bacteroides thetaiotaomicron*^44^ on EggNog v5.0 (online)^87^. The orthologous group was identified as COG0156 (Amino acid transport and metabolism: 8-amino-7-oxononanoate synthase activity) and abundance was acquired using a python script called geneshot_extract_gene_abund.py (https://github.com/Golob-Minot/geneshot/wiki/Extracting-the-abundances-for-a-single-gene) on the geneshot results of the metagenomic data.^37^ *Bacteroides fragilis* serine palmitoyltransferase (encoded by GenBank# EXZ60402.1) was queried among the *B. xylanisolvens* reference strains found in our metagenomic sequencing. *B. xylanisolvens* strains were subjected to NCBI BLASTP (http://blast.ncbi.nlm.nih.gov) using the default parameters. The top hits of each sample are listed with scores (bits) and E-values in Table S10. Samples with scores > 500 and E-values less than E-50 were considered a hit, i.e., containing a close homology to the query.

### Fecal lipid quantification

Lipidomics was performed on human fecal samples using 50-100 mg of feces (Northwest Metabolomics Research Center at the University of Washington). Feces were heated at 70°C for 8 hours in a dry oven. Samples were then homogenized in a fixed volume of water and volume equivalent to 10 mg was used. Lipidomics was performed with the Sciex Lipidyzer platform as described previously.^95^ Data was plotted using log2 fold change of average concentrations of lipid species (µM) of OCV responders (N=8) and nonresponders (N=8). Log10 (p-value) which was obtained using multiple Mann-Whitney test.

### Bacterial strains, growth conditions, and identification

*Bacteroides xylanisolvens* was isolated from human fecal samples from the study participants^26^ using Brain-Heart Infusion (BHI) agar (BD) supplemented with 5% sheep’s blood (Hemostat Labs) and Laked Brucella Blood agar with Kanamycin and Vancomycin (LKV) plates (Anaerobe Systems). *Bacteroides koreensis* was isolated using Gifu Anaerobic Media (HiMedia) supplemented with porcine hemin (10 mg/L; MP Biomedicals) and menadione (5 mg/L; MP Biomedicals) and LKV plates. Bacterial identity was confirmed using ∼1000 bp Sanger sequencing of the 16S rRNA gene with >97% identity. Liquid cultures of *Bacteroides* strains were prepared in BHI broth (BD Diagnostic Systems) supplemented with 0.005 mg/mL porcine hemin, 0.5 g/L L-cysteine (Sigma), and 0.05 ng/mL Vitamin K1 (Sigma) and cultivated at 37°C in a vinyl anaerobic chamber (Coy) with <50 ppm O_2_. Glycerol stocks were prepared using BHI-supplemented media 20% glycerol and stored in −80°C ultra-low freezers. *Bacteroides* strains were maintained on Tryptic Soy Agar with 5% sheep’s blood plates and grown in BHI supplemented media for 24-48 hours. The *V. cholerae* strain JBK70 was gifted by Dr. Jim Kaper.^45^ *V. cholerae* was maintained in lysogeny broth (LB) or LB agar at 37°C in aerobic conditions.

### Mammalian tissue culture cell line and maintenance

THP-1 cells (ATCC) were maintained in RPMI-1640 media (containing L-glutamine and 25 mM HEPES) (Corning) supplemented with 10% fetal bovine serum (Fisher) and penicillin and streptomycin (Gibco) in 5% CO_2_ and 37°C conditions.

### Bligh-Dyer Lipid extraction and Mild Alkaline Hydrolysis

To inhibit *spt* and de novo sphingolipid production, 5 µM myriocin (Cayman Chemicals) was added to bacterial culture. Forty-eight hour liquid bacterial cultures in BHI-supplemented with or without myriocin with OD > 1.00 were used for lipid extraction. Bacterial cultures were normalized to an optical density (OD 600 nm) of 1-1.5 using sterile phosphate-buffered saline (PBS; Corning) with total volume of 5 mL. Samples were then centrifuged at 4,000 x g for 20 minutes at 4°C. The supernatant was removed, and the bacterial pellets were resuspended in sterile PBS and transferred to 1.5 mL microtubes and washed twice using PBS. Bacterial pellets were then resuspended in 100 μL sterile water and treated with 400 μL chilled 1:2 parts chloroform:methanol (Fisher Chemical). Samples were vortexed for 5 minutes at max speed using a vortex microtube adaptor. Samples were then treated with 100 µL chilled chloroform and 100 μL sterile molecular-grade water (Corning) and vortexed for 1 minute. Samples were then centrifuged at 2,000 x g for 10 minutes. The bottom layer containing chloroform and lipids were collected into a new tube and air-dried in a sterile environment. Samples were stored at −80°C until resuspended in 500 μL of cell culture media corresponding to the mammalian cells used for the assay. Reconstituted lipids were stored at −20°C. Mild alkaline hydrolysis for removal of phospholipids from lipid extracts was performed as previously described.^46^ Briefly, the lipid extracts before drying were treated with 0.02N sodium hydroxide (NaOH; Acros Organics) for 30 minutes at 37°C. Lipids were then dried, stored, and reconstituted as above.

### THP-1 cell differentiation and treatment with bacterial lipids and JBK70

THP-1 cells were treated with 50 ng/mL PMA (phorbol 12-myristate 13-acetate) (Invitrogen) and plated into 24-well tissue-culture treated plates using 5×10^5^ cells/well. Cells were incubated for 48 hours prior to exposure to additional stimuli. Adherent cells were rinsed using sterile PBS and the cell media was replaced. Bacterial lipids were applied to differentiated THP-1 cells in 1:100 concentrations for 24 hours for pre-conditioning. Cells were rinsed with sterile PBS and then treated with heat-killed JBK70 (1:10) for a final dilution of 1:100. After 18 hours, the supernatant of cells was collected.

### Preparation of Heat-Killed JBK70

*V. cholerae* strain JBK70 was inoculated into LB broth from a glycerol stock and incubated at 37°C with agitation for 24 hours. Opaque cultures were centrifuged and resuspended in equal volume (4 mL) of cell culture media corresponding to the mammalian cells used for the assay. Samples were then diluted 1:10 in cell media and heated at 95°C for 30 minutes. Heat-treated samples were cooled to room temperature before application to mammalian cells. Aliquots of heat-killed JKB70 were plated on LB agar to ensure no live bacteria was present.

### Enzyme linked immunosorbent assays (ELISA) and cytokine measurements

Cell supernatants were analyzed using ELISA for cytokine production. ELISA kits were purchased from R&D systems and used according to manufacturer’s instructions, including IL-1*β*, IL-6, IL-10, TNF-*α*, MCP-1, and IL-8. For multiplex assays, samples were analyzed using the human cytokine proinflammatory focused 15-Plex Discovery Assay (HDF15) (Eve Technologies Corporation, Calgary, Canada). Briefly, samples stored at −80°C were thawed and centrifuged at 3,000 x g for 10 minutes at 4°C. The supernatants of samples were collected and shipped to Eve Technologies for processing.

### RNA extraction and sequencing

THP-1 cells were collected after exposure to *V. cholerae* and/or bacterial sphingolipids for RNA sequencing. After collecting the cell supernatant for cytokine analysis, adherent cells were treated with TRIzol (Invitrogen) and transferred into a 1.5 mL microtube. Chloroform was added, centrifuged at 4°C, and the aqueous layer was transferred into a new 1.5 mL tube and treated with 70% ethanol (Spectrum). Samples were then processed using the Qiagen RNeasy kit (Qiagen) according to manufacturer’s instructions with on-column DNase treatment. RNA was eluted using 30 µL molecular-grade water and additionally treated with TURBO DNase (Ambion). RNA quality was assessed using TapeStation (Agilent). RNA was prepared for sequencing using Stranded Total RNA Prep with Ligation with Ribo-Zero Plus (Illumina) and RNA UD Indexes Set A, Ligation (Illumina). Sequencing was performed using P3 Reagents (Illumina) on a NextSeq 2000 as pair-ended reads.

### RNAseq preprocessing and analysis

Fastq files were obtained and RNA reads were curated and aligned using the RNA snakemake pipeline SEAsnake.^96^ Briefly, RNA quality was assessed with FastQC, adapters were removed, and low quality sequences were filtered using AdapterRemoval. Reads were aligned to the Homo sapiens genome GRCh38 (release 108) using STAR. Alignments were quality checked and filtered using samtools and Picard and read counts were determined using Subread. RNA-samples were then quality assessed and removed if libraries had <1,000,000 sequences, greater than 1 coefficient of variation (CV) coverage, and less than 75% alignment to genome. No sample or library was removed using these criteria. Low abundance genes with a minimum CPU of 0.1 and present in a minimum of 2 samples were filtered out. 4244 (21.22%) of 20001 genes were removed. Gene counts were then normalized by trimmed mean of M (TTM) and voom (log2 transformation of counts per million (CPM)). Analysis was then performed on the following comparisons: SL-containing with and without HK JBK70, SL-free with and without HK JBK70, SL-containing and SL-free without HK JBK70, and SL-containing and SL-free with HK JBK70. Linear modeling was performed using kmFit (Kimma software on R: https://github.com/BIGslu/kimma)^97^ with weighted factors to obtain the estimate (log2 fold change) and FDR values. Genes were considered significant with FDR < 0.3 and log2FC greater than 1 or less than −1. Enrichment pathways were determined using the R package BIGslu/SEARchways (https://github.com/BIGslu/SEARchways) with BIGprofiler that employs clusterProfiler using the gene set data base and hallmark gene sets available in the Broad Molecular Signatures Database (MSigDB).^98^ Pathways were considered significant with FDR < 0.05.

### Statistical Analysis

Visualizations, figures, and statistical testing were generated using R (v4.0.2) and Graphpad Prism (v10) unless otherwise specified. Strain trees were generated using GToTree (v1.5.51)^99^ and visualized using the Interactive Tree of Life (v6.1.2)^100^. Shannon indices were compared between sexes using a two-tailed Mann Whitney U test. A Kruskal-Wallis test with Dunn’s adjustment for multiple comparisons was used to assess differences in Shannon index between age groups as well as between responders and nonresponders for each vaccine response measures.

### Ethics approval and consent for publication

This study was approved by the Research Review and Ethics Review Committee of the icddr,b, in Dhaka, Bangladesh. Approvals for this work were also granted by the Institutional Review Board of the Massachusetts General Hospital and the University of Washington. Written informed consent was obtained for all participants in this study. For children below 17 years of age, permission of at least one parent/guardian was required for participation. For those aged 11-17 years of age, participant written assent was also obtained.

## Acknowledgments

We would like to thank Dr. Stephen J. Salipante, Dr. Kim Dill-McFarland, and the Microbial Interactions & Microbiome Center (mim_c) at the University of Washington for help and support for the RNA sequencing; Dr. Tom Hawn for donating THP-1 cells; Dr. Jim Kaper for the *V. cholerae* JBK70 strain. We would like to thank Meti Debela, Chelsea N. Dunmire, Dr. Tsoni Peled, Dr. Libin Xu, Emily Pruitt, the study participants, and the icddr,b laboratory, field, and data management staff. The icddr,b gratefully acknowledges the government of the People’s Republic of Bangladesh, and Canada. The funding bodies had no role in the design of the study, data collection, analysis, interpretation of data or writing of the manuscript.

This work was supported by the NIH [NIAID R01 AI103055 to J.B.H. and R.C.L.; R01 AI099243 to J.B.H. and F.Q, K08 AI123494 to A.A.W.; R01 106878 to E.T.R, and NIGMS R35 GM133420 (support of S.S.M, PI: Amy D. Willis); T32HD007233 to D.C.], Fogarty International Center [D43 TW005572 to T.R.B, and K43 TW010362 to T.R.B], the University of Washington [Chief of Medicine Award and Royalty Research Foundation support to A.A.W.], the Government of the People’s Republic of Bangladesh (to the International Centre for Diarrheal Disease Research [icddr,b]), Global Affairs Canada (to the icddr,b),.

## Author’s Contributions

D.C. contributed to methodology, formal analysis and investigation, data curation, visualization and drafting of the original manuscript. F.J.H. contributed to methodology, code development, formal analysis and investigation, visualization, and drafting of the original manuscript. H.A.B., A.A., and M.H.K. contributed to the investigation and drafting of the original manuscript. F.C., T.R.B., A.I.K., P.C.K., and P.D. contributed to the investigation, data curation, and editing of the manuscript. M.G.D., S.M.M., and A.R., contributed to the experimental investigation. R.C.L, E.T.R., and J.B.H. contributed to the conceptualization of the project and editing of the manuscript. F.Q. contributed to the conceptualization of the project, investigation, data curation, and editing of the manuscript. S.S.M. contributed to the methodology, code development, formal analysis, and drafting/editing of the original manuscript. A.A.W. contributed to the conceptualization of the project, formal analysis, and drafting/editing of the original manuscript.

