## Supplemental Figures for "Gut bacteria-derived sphingolipids alter innate immune responses to oral cholera vaccine antigens"

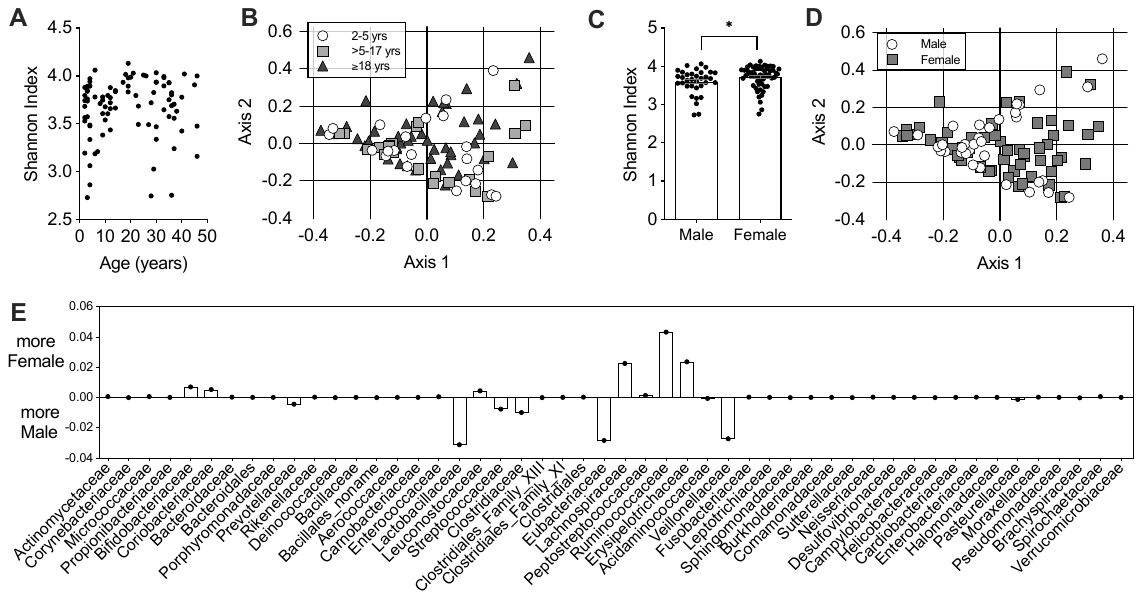

### Figure S1. Diversity by age and sex.

(**A**) Shannon Index by age. No difference was observed between participants by Kruskal-Wallis testing when samples were divided by age group (2-5, >5-17, ≥18 years of age, p>0.05). (**B**) Beta diversity between age groups measured by Bray-Curtis Dissimilarity, Principal Component Analysis (PCA). No difference was observed in statistical testing between age groups via Kruskal-Wallis testing (p>0.05). (**C**) Shannon diversity according to participant sex. Bars represent mean with SD, and male and female participants were found to have a significantly different Shannon Index by Mann-Whitney U testing (P=0.03). (**D**) Beta diversity between sexes measured by Bray-Curtis Dissimilarity. No difference was found between sexes by Mann-Whitney U testing (p>0.05). Dots in each graph represent one participant. (**E**) Abundance differences in baseline gut microbiota populations by sex (family level shown). Positive values on the X axis represent a greater proportion of gut microbes in that family in female participants. All families with abundance <1% in this study population are shown.

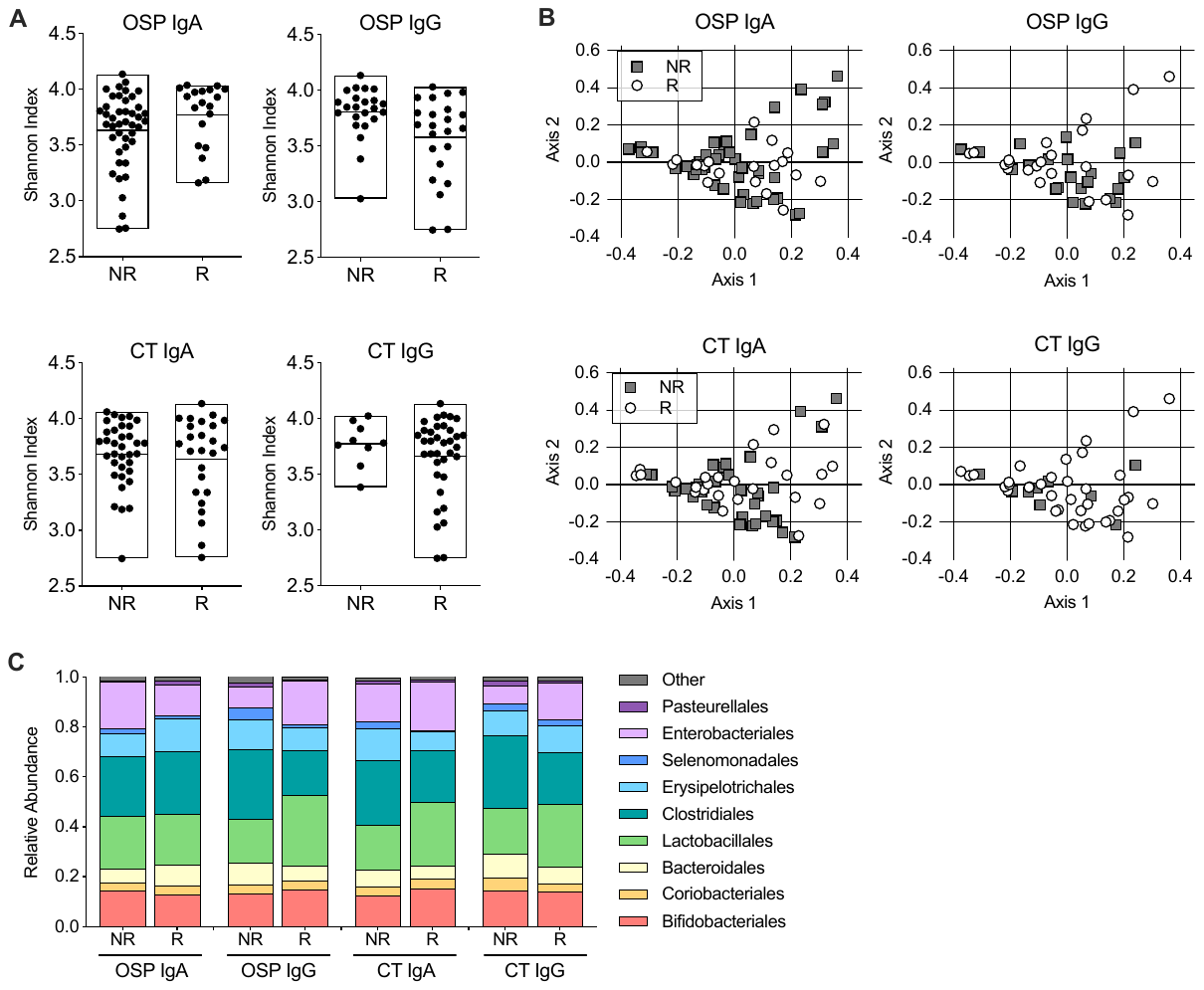

### Figure S2. Microbial diversity by memory B cell response.

(**A**) Shannon index of vaccine responders (R) and nonresponders (NR) by MBC-specific responses. Box represents max and min with middle line at the mean. No difference was found between groups via Mann-Whitney U testing (p>0.05 in each group). (**B**) PCA demonstrating beta diversity between R and NR measured by Bray-Curtis Dissimilarity within each vaccine response measure. No difference was seen between R and NR groups by Mann-Whitney U testing (p>0.05). Dots in each graph represent one participant.

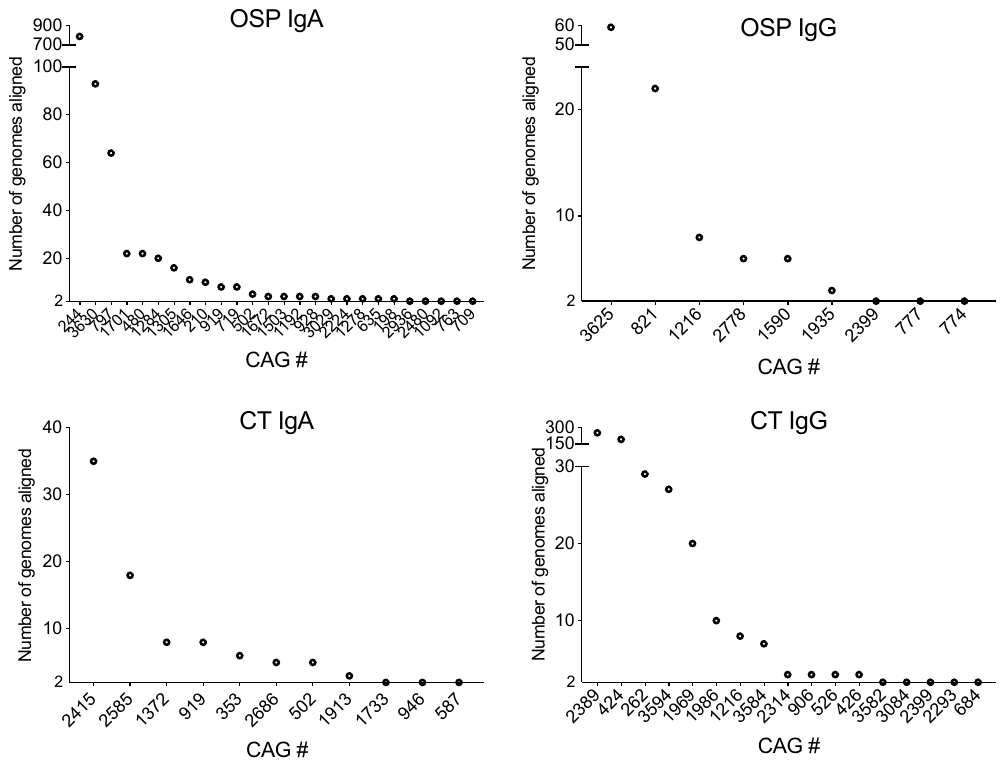

### Figure S3. Number of genomes aligning to significant CAGs found to correlate with OSP- and CT-specific IgA and IgG producing MBCs after OCV.

Each circle represents one significant CAG containing ≥2 genes with q≤0.1 for an association with the listed vaccine response measure. The Y axis represents the number of reference genomes to which genes in those CAGs align with >90% sequence identity. Significant CAGs aligning to only one bacterial genome are not shown (20 CAGs for OSP IgA, 3 CAGs for OSP IgG, 12 CAGs for CT IgA, 5 CAGs for CT IgG).

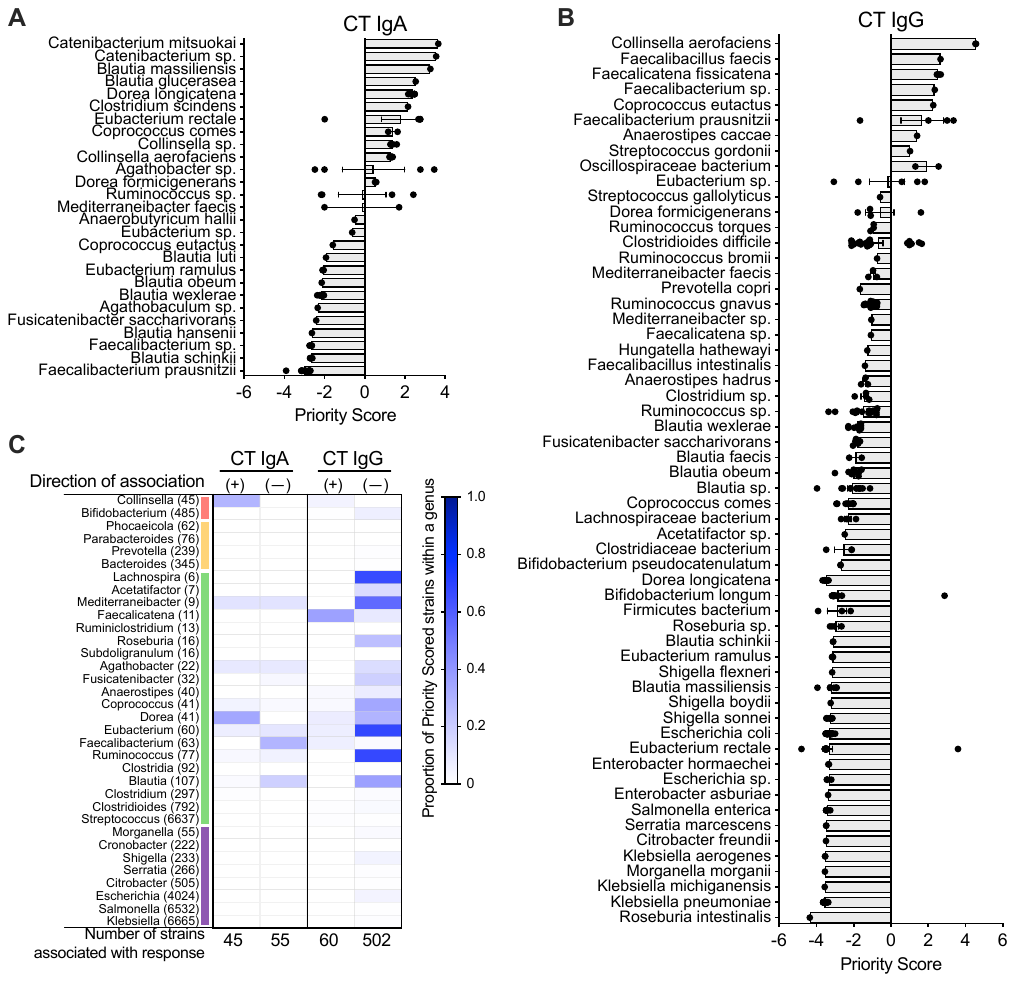

### Figure S4. Top strains associated with CT-specific MBC responses after OCV.

Species containing strains with a priority score greater than 0.5 are shown for (**A**) CT IgA and (**B**) CT IgG MBC responses. Each dot indicates the priority score of a specific strain within the species listed on the left in each row. For each species, bars represent mean with SEM of priority scored strains. Full strain data and priority scores for each MBC response are shown in Tables S3-6.

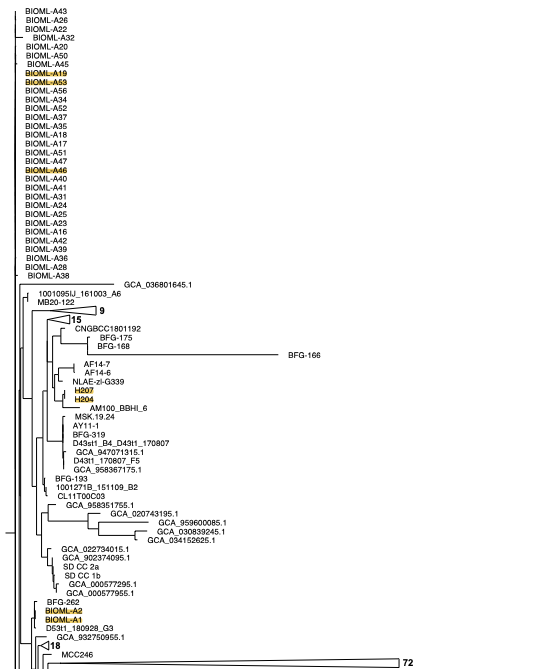

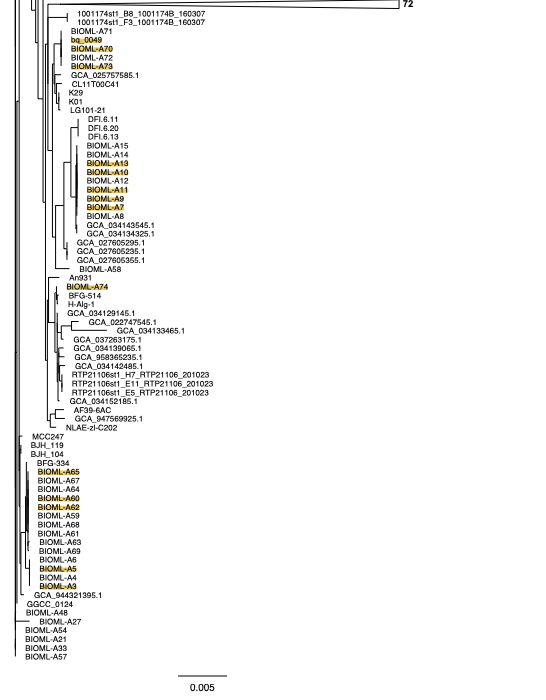

### Figure S5. Phylogenetic tree of *Bacteroides xylanisolvens* strains.

*Bacteroides xylanisolvens* NCBI genomes are shown, and Bx strains from the stool of our study population are highlighted. Tree assembly was performed using GToTree and drawn with FigTree.

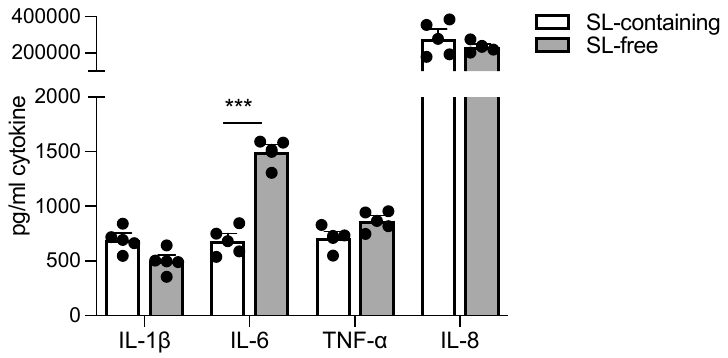

### Figure S6. Stimulation of human macrophages with *Bacteroides xylanisolvens* sphingolipid-free lysate induces increased IL-6 responses.

Cytokine response was measured in supernatant of THP-1 derived macrophages incubated with *Bacteroides xylanisolvens* lysate that was grown with or without myriocin, SL-free and SL-containing, respectively for 18 hours. Statistical analysis was performed using multiple unpaired t tests with FDR two-stage set-up method of Benjamini, Kriege and Yekutieli. ***, P≤0.001. Bars indicate mean with SEM.

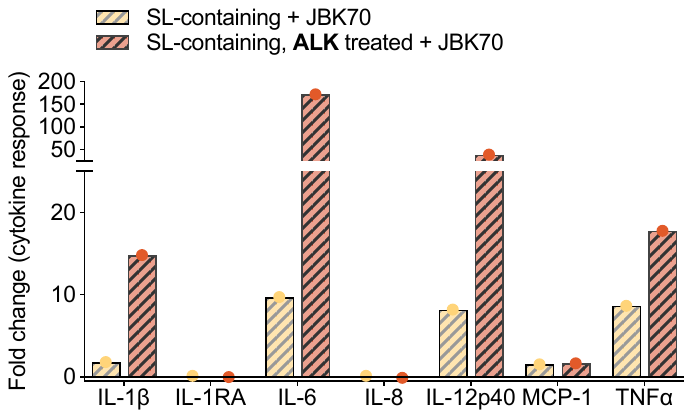

### Figure S7. Cytokine response of THP-1 derived macrophages to heat killed JBK70 after preconditioning with SL-containing lipids with and without phospholipids.

THP-1 derived macrophage fold change cytokine response to HK JBK70 after preconditioning with *B. xylanisolvens* lipids with or without a mild alkaline hydrolysis treatment (ALK) that eliminates non-sphingolipid phospholipids demonstrated increases in response compared to SL-containing fractions alone. Cytokines were measured using a 15-cytokine multiplex assay.

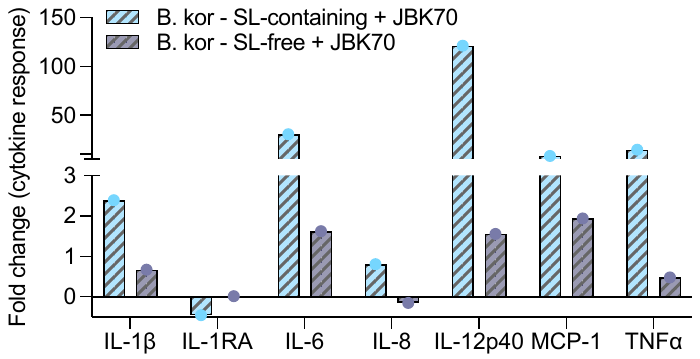

### Figure S8. Cytokine response of THP-1 derived macrophages to heat killed JBK70 after preconditioning with *Bacteroides koreensis* lipids.

THP-1 derived macrophage fold change cytokine response to HK JBK70 after preconditioning with *Bacteroides koreensis* SL-containing and SL-free lipid fractions. Cytokines were measured using a 15-cytokine multiplex assay.

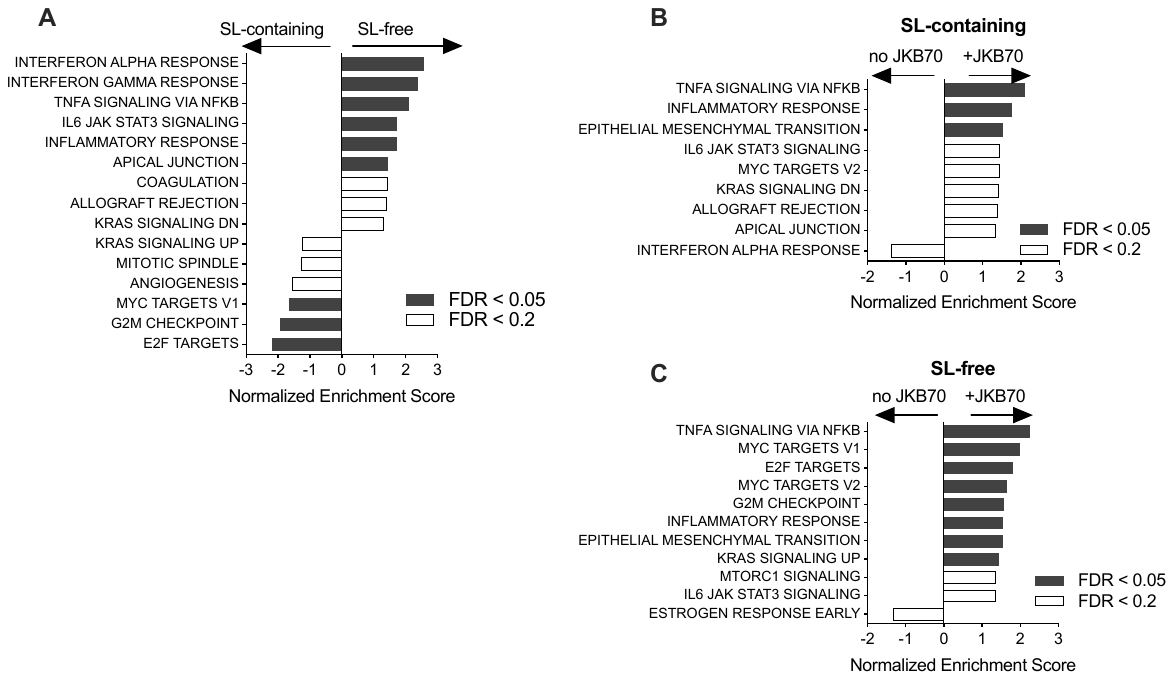

### Figure S9. Innate immune pathway activation in human macrophages exposed to JBK70 was greater after preconditioning with Bx SL-containing fractions compared to SL-free fractions.

Pathway analysis and gene categories generated using the gene set enrichment analysis method on fold change values of all genes present in the dataset is shown. Pathways enriched between (**A**) SL-containing and SL-free preconditioning without JBK70 stimulation, (**B**) SL-containing fraction with and without JBK70 stimulation, (**C**) and SL-free with and without JBK70 stimulation.

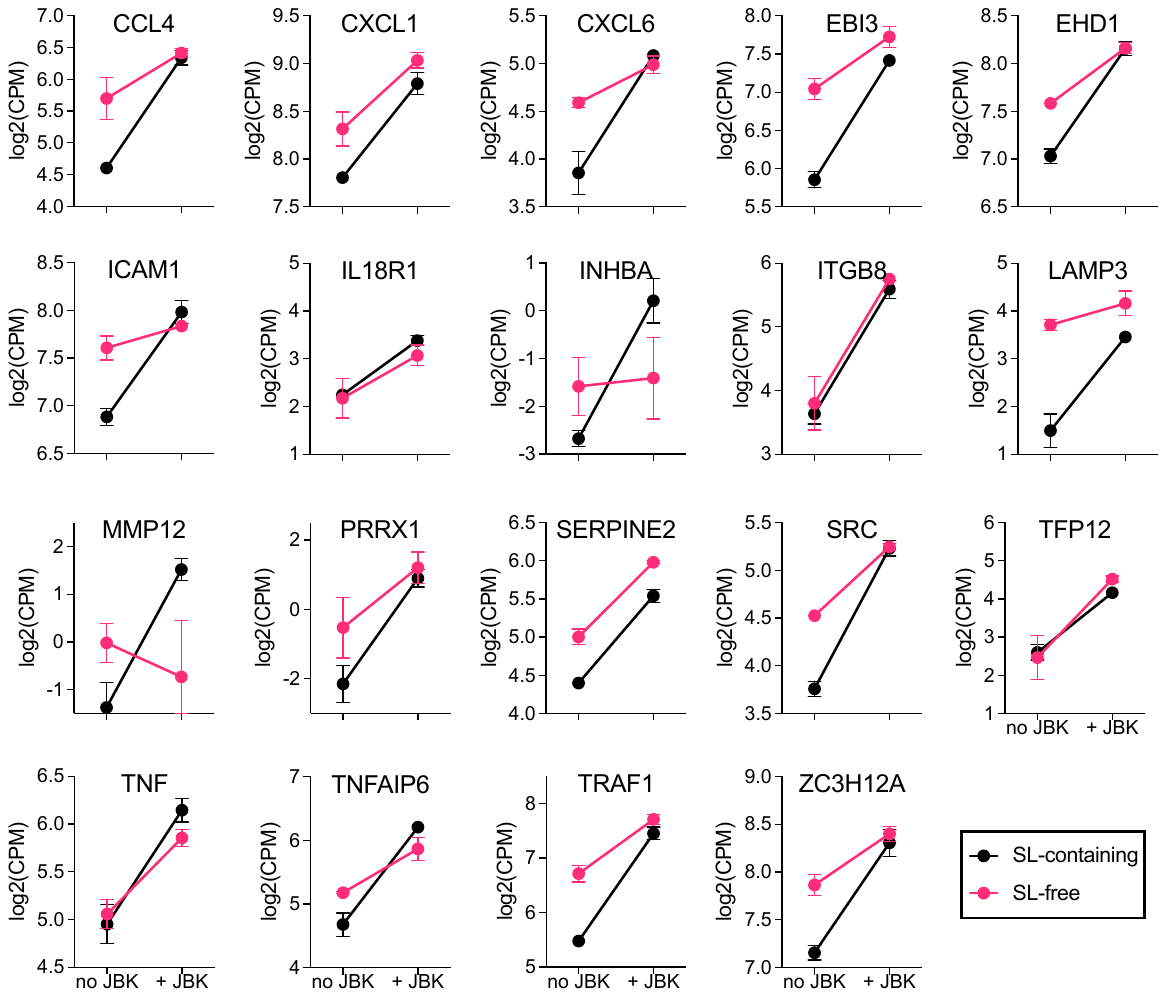

### Figure S10. Differential gene expression in human macrophages after preconditioning with Bx SL-containing or SL-free lipid fractions and stimulation with JBK70.

Gene expression is shown as log2 counts per million (CPM) of the 19 genes in the enriched pathways identified using gene set enrichment analysis, described in Figure 5C-D.
